# Honey bee suppresses the parasitic mite Vitellogenin by antimicrobial peptide

**DOI:** 10.1101/2020.02.29.970913

**Authors:** Yunfei Wu, Qiushi Liu, Benjamin Weiss, Martin Kaltenpoth, Tatsuhiko Kadowaki

## Abstract

The negative effects of honey bee parasitic mites and deformed wing virus (DWV) on honey bee and colony health have been well characterized. However, the relationship between DWV and mites, particularly viral replication inside the mites, remains unclear. Furthermore, the physiological outcomes of honey bee immune responses stimulated by DWV and the mite to the host (honey bee) and perhaps the pathogen/parasite (DWV/mite) are not yet understood. To answer these questions, we studied the tripartite interactions between the honey bee, *Tropilaelaps mercedesae*, and DWV as the model. *T. mercedesae* functioned as a vector for DWV without supporting active viral replication. Thus, DWV negligibly affected mite fitness. Mite infestation induced mRNA expression of antimicrobial peptides (AMPs), Defensin-1 and Hymenoptaecin, which correlated with DWV copy number in honey bee pupae and mite feeding, respectively. Feeding *T. mercedesae* with fruit fly S2 cells heterologously expressing honey bee Hymenoptaecin significantly downregulated mite *Vitellogenin* expression, indicating that the honey bee AMP manipulates mite reproduction upon feeding on bee. Our results provide insights into the mechanism of DWV transmission by the honey bee parasitic mite to the host, and the novel role of AMP in defending against mite infestation.

## Introduction

Large-scale loss of managed honey bee colonies has been recently reported across the globe (Goulson *et al.*, 2015). Since pollination by honey bees is vital for maintaining ecosystems and the production of many crops (Klein *et al.*, 2007; Aizen and Harder, 2009), prevention of honey bee colony losses has become a major focus in both apiculture and agriculture. Colony losses have often been associated with the ectoparasitic mites *Varroa destructor* and *Tropilaelaps mercedesae*, which feed on honey bees and transmit honey bee viruses, particularly deformed wing virus (DWV) to the host (de Miranda and Genersch, 2010; Rosenkranz et al., 2010; Chantawannakul et al., 2018). In the absence of mites, DWV copy numbers remain low in honey bees without specific symptoms (covert infection). However, DWV levels associated with honey bees are dramatically increased in mite infested colonies (Shen *et al.*, 2005; Khongphinitbunjong *et al.*, 2015; Forsgren *et al.*, 2009; Wu, Dong and Kadowaki, 2017). These honey bees often show multiple symptoms (overt infection), which include the death of pupae, deformed wings, shortened abdomen, and reduced lifespan (Yue and Genersch, 2005; Tentcheva et al., 2006; de Miranda and Genersch, 2010; Rosenkranz et al., 2010). Winter colony loss is strongly correlated with the presence of DWV and *V. destructor* (Highfield *et al.*, 2009; Nazzi and Le Conte, 2016).

Although the impacts of DWV and *V. destructor* on individual honey bees and colonies are well characterized, the actual relationship between DWV and honey bee mite is not yet understood. Several studies have suggested that DWV replicates in *V. destructor* and that more virulent DWV strains are amplified for transmission to honey bees (Martin *et al.*, 2012; Ryabov *et al.*, 2014). However, the results of other studies are inconsistent with this view (Erban *et al.*, 2015; Dong *et al.*, 2017; Posada-Florez *et al.*, 2019). Thus, it is important to address this issue to uncover the mechanism by which mites function as vectors for DWV.

DWV copy numbers in mites can exceed 10^6^ (Wu, Dong and Kadowaki, 2017; Posada-Florez *et al.*, 2019). Thus, DWV could have significant effects on mite physiology. Previous studies have reported that DWV infection and/or *V. destructor* infestation induce honey bee immune responses that include synthesis of antimicrobial peptides (AMPs) (Gregorc *et al.*, 2012; Kuster, Boncristiani and Rueppell, 2014). However, their effects on the host (honey bee), pathogen (DWV), and parasite (mite) are still uncharacterized. AMPs were originally identified as short positively-charged peptides that inhibit the viability of bacteria and fungi (Bahar and Ren, 2013; Hanson and Lemaitre, 2019). Since AMPs are induced under various conditions, their physiological functions could be more diverse and remain to be tested.

In this study, we first examined whether *T. mercedesae* functions as a bona fide vector for DWV, and then characterized the effects of DWV on mites to understand the precise relationship. We also examined the immune responses of honey bee pupae to *T. mercedesae* infestation and found that Hymenoptaecin down-regulates the mite *Vitellogenin* (*Vg*) gene. Using these results, we discuss the mechanisms of DWV transmission by the honey bee parasitic mites, as well as how the host (honey bee) defends against the parasite (mite) by suppressing reproduction.

## Results

### Role of *T. mercedesae* as a vector for DWV

To directly test whether *T. mercedesae* functions as a vector to transmit DWV to honey bees, we collected 15 pupae with white eyes from a mite-free colony and added them individually to gelatin capsules for seven days together with single *T. mercedesae* sampled from another mite infested colony. As a control, we incubated 18 pupae without mites under the same conditions. The test pupae (the ones artificially infested by the mites) contained higher DWV copy numbers than the control pupae (Fig. 1A). We also found a positive correlation between DWV copy number in pupae and infesting mites. The pupa-mite pairs were separated into two groups with low and high DWV loads (Fig. 1B). Sanger sequencing of the PCR amplicons (Supplementary Fig. 1) revealed that ten pupa-mite pairs were infected by a single variant of DWV. Two pairs were infected by multiple variants in both pupae and mites. Two pairs were infected by single and multiple variants in pupae and mites, respectively. One pair was infected by single and multiple variants in mite and pupa, respectively (Supplementary Table 1). A phylogenetic tree was constructed using DWV sequences obtained from test pupae, infesting mites, as well as the control pupae. In case multiple DWV variants are present in pupae and mites, only samples in which we could identify the dominant variants were analyzed. All 18 control pupae were infected by multiple variants at very low level and the eight samples (Bee-C2, 6, 9, 10, 11, 14, 15, and 16) contained the dominant DWV variant, which was identical to the one present in seven test pupae (Bee-1, 4, 7, 8, 9, 12, and 14) and five infesting mites (Mite-1, 7, 8, 9, and 12). Three pupa-mite pairs (Bee/Mite-3, 13, and 15) contained the same variant. Mite-4 and Mite-14 shared the same variant, which was different from the one present in the infested pupae (Bee-4 and Bee-14). These two pairs represent the example that pupa and the infesting mite do not share the same DWV variant. As shown in Figure 1C, six pupa-mite pairs (Bee/Mite-3, 5, 10, 11, 13, and 15) out of 15 pairs were infected by four variants that were different from the ones present in the control pupae (Fisher’s exact test; *P* < 0.005). These results demonstrated that these variants were transferred from the infesting mites derived from a colony that was different from the one in which all pupae were sampled. To test the possibility that *T. mercedesae* also functions as a mechanical vector for DWV, we induced wounds at the pupal thorax with a sterile needle and compared the copy numbers of DWV in the heads of wounded and control (untreated) pupae. The copy numbers were higher in the wounded pupae than in the control pupae (Fig. 1D), suggesting that wound induction alone is sufficient to stimulate replication of the endogenous DWV in honey bee pupae.

**Figure 1.**
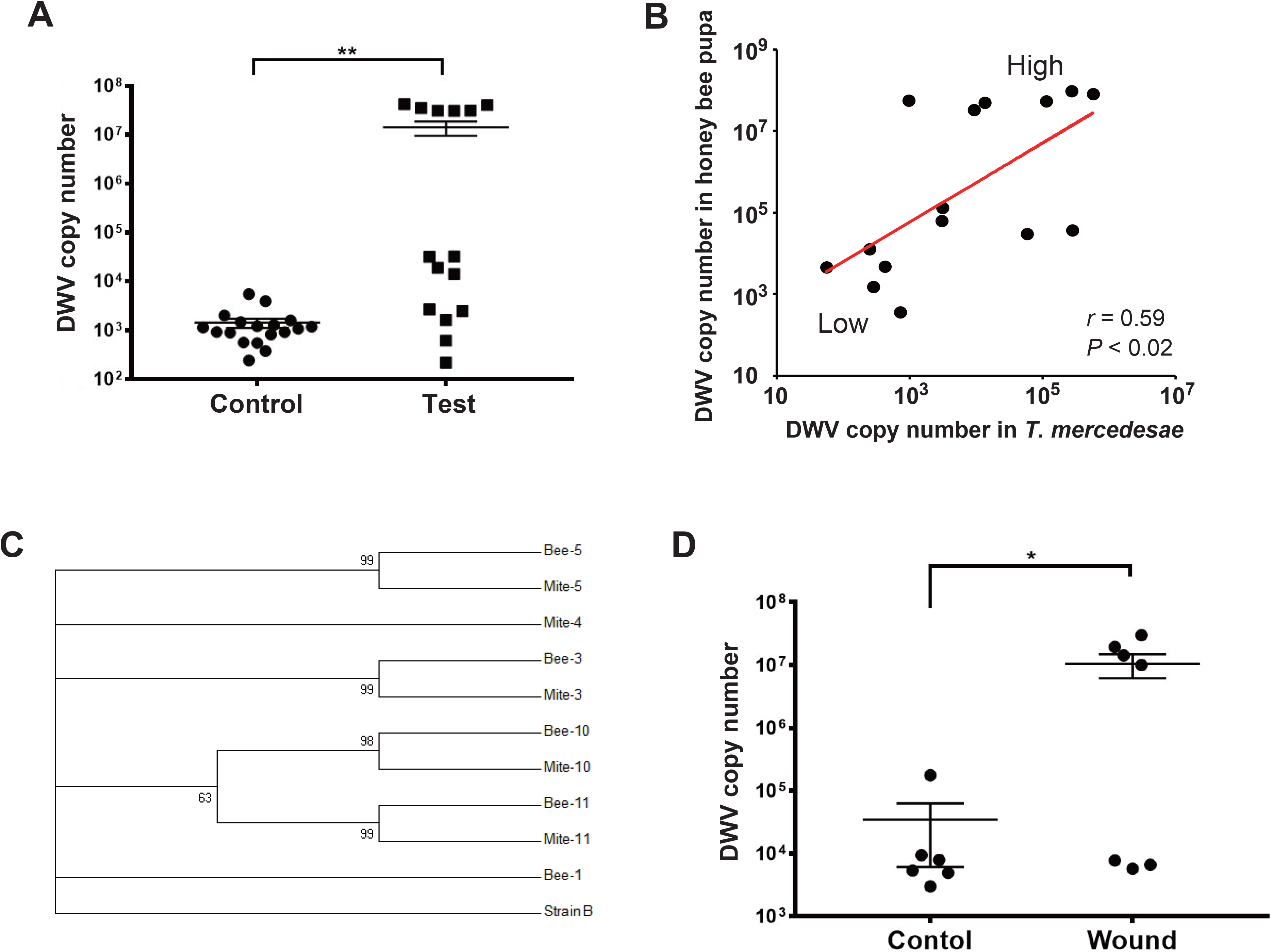
*T. mercedesae* functions as a vector for DWV. (A) DWV copy numbers in honey bee pupae artificially infested by *T. mercedesae* (Test, n =15) and the ones without mites (Control, n = 18). Mean values with error bars (± SEM) are indicated. Two groups are statistically different by Brunner-Munzel test (**, *P* < 1.3E-06). (B) DWV copy numbers in individual honey bee pupae and the infesting *T. mercedesae* are plotted on the Y- and X-axis, respectively. The red line represents a fitted curve and there are pupa/mite pairs with high or low DWV copy numbers. The Pearson correlation value and *P*-value are also shown. (C) Maximum-likelihood condensed phylogeny of DWV isolates from honey bee pupae (Bee) and *T. mercedesae* (Mite) was constructed based on partial VP2 and VP1, and full VP4 sequences. In case the same variant is shared by multiple samples, only one representative is indicated. Bootstrap values are shown at the corresponding node of each branch and DWV type B strain was used as an outgroup. (D) DWV copy numbers in wounded (Wound, n = 7) and untreated (Control, n = 6) honey bee pupae. Mean values with error bars (± SEM) are indicated. Two groups are statistically different by Brunner-Munzel test (*, *P* < 0.041).

### Impact of DWV on *T. mercedesae*

To examine the effects of DWV on *T. mercedesae*, we collected mites carrying either high or low DWV copy numbers by RT-PCR, pooled ten mites into one group, and then compared the gene expression profiles of replicates using RNA-seq. The average mapping rates of RNA-seq reads derived from the mites with high and low DWV loads to *T. mercedesae* genomes were 38.7 % and 83.0 %, respectively, and those to the DWV genome were 22.8 % and 0.05 %, respectively (Supplementary Table 2). We identified a few differentially expressed genes between mites with low and high DWV loads (Supplementary Table 3). The high DWV load had little effect on the transcriptome profile of *T. mercedesae*, suggesting that DWV does not actively infect/replicate the mite cells. To test this hypothesis, we analyzed the protein extracts of individual honey bee pupal heads and *Tropilaelaps* mites to detect VP1 (capsid protein, structural protein) and RdRP (RNA dependent RNA polymerase, non-structural protein) of DWV. As shown in Figure 2A, two bands of 90 and 53 kDa were specifically detected by anti-RdRP antibody, suggesting that the large and small bands represent the precursor of RdRP fused with 3C-protease (3C-Pro) and the matured RdRP, respectively. The cleavage between 3C-Pro and RdRP appears to be rate-limiting for DWV, similar to other picornaviruses (Jiang *et al.*, 2014). RdRP and the precursor were detected in the VP1-positive, but not VP1-negative, honey bee pupal head. However, these bands were not detected in the *Tropilaelaps* mite, irrespective of the presence of VP1. These results indicated that DWV infects and actively replicates in the honey bee pupal head, but not the mite cells. Lack of active synthesis of DWV protein in *Tropilaelaps* mites was also supported by their large differences in codon usage. The codon usage of DWV is apparently adapted to that of the original host, *A. mellifera*, as previously reported (Chantawannakul and Cutler, 2008), but quite different from that of mite (*r* = 0.046, *P* < 0.73) (Fig. 2B and C). To uncover the major sites for localization of DWV in *Tropilaelaps* mite, transverse thin sections of mites were immunostained using anti-VP1 antibody. DWV was primarily localized in the lumen of the entire midgut as large dense spheres (Fig. 3), consistent with the lack of active viral replication. There was no specific staining with the pre-immune serum (Supplementary Fig. 2).

**Figure 2.**
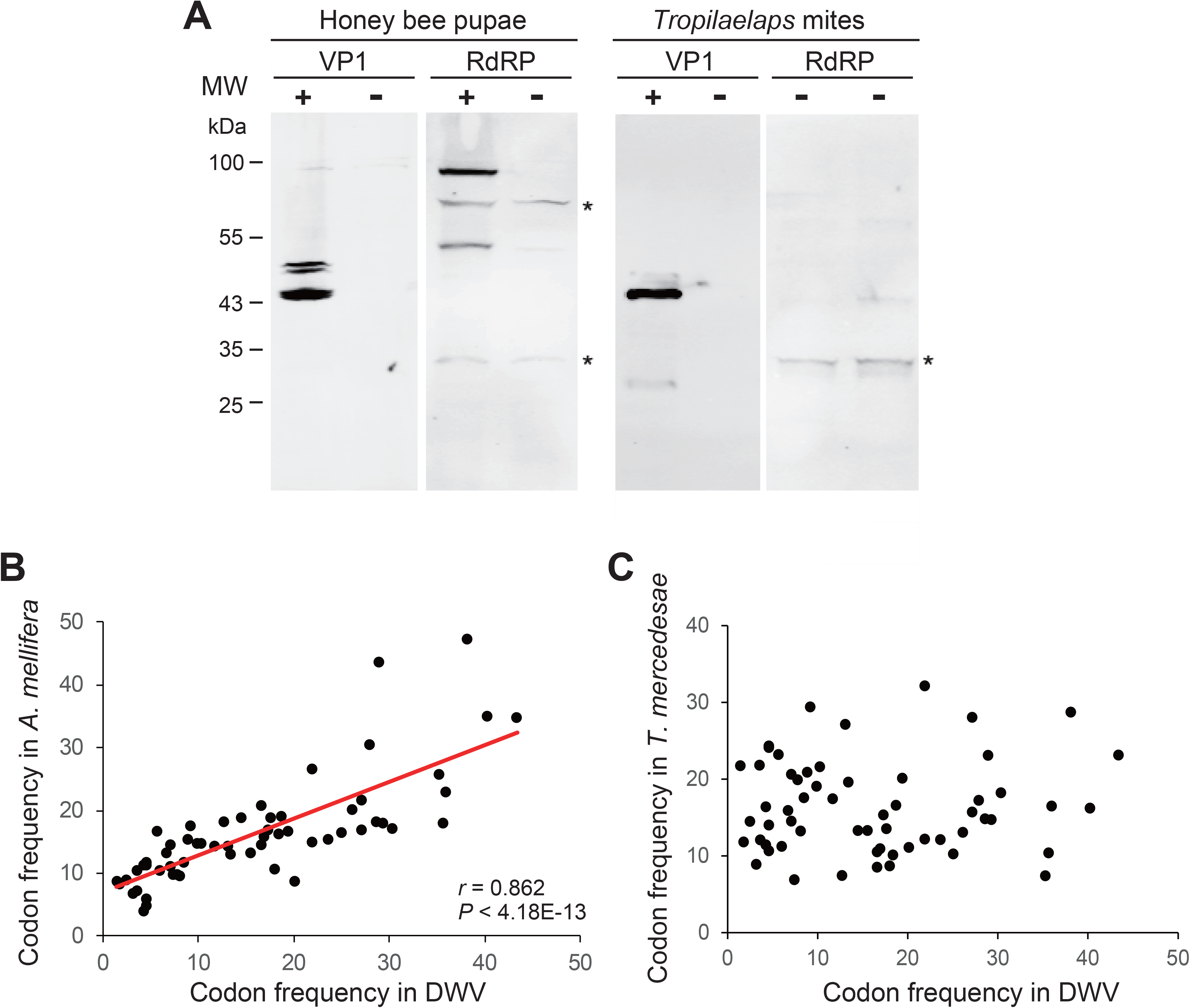
Lack of active replication of DWV in *T. mercedesae*. (A) Detection of DWV VP1 and RdRP in the lysates of individual honey bee pupae and *Tropilaelaps* mites by western blot. A major 47 kDa band was detected in VP1-posiitive (+) but not negative (−) pupa and mite; however, RdRP (90 and 53 kDa bands) was only present in the VP1 positive pupa. The asterisks represent non-specific proteins cross reacted with anti-RdRP antibody. The size of protein molecular weight marker (MW) is at the left. (B) Frequencies of each codon out of 1,000 codons in DWV and *A. mellifera* genomes are plotted on the X- and Y-axis, respectively. The red line represents a fitted curve and the Pearson correlation value and *P*-value are also shown. (C) Frequencies of each codon out of 1,000 codons in DWV and *T. mercedesae* genomes are plotted on the X- and Y-axis, respectively.

**Figure 3.**
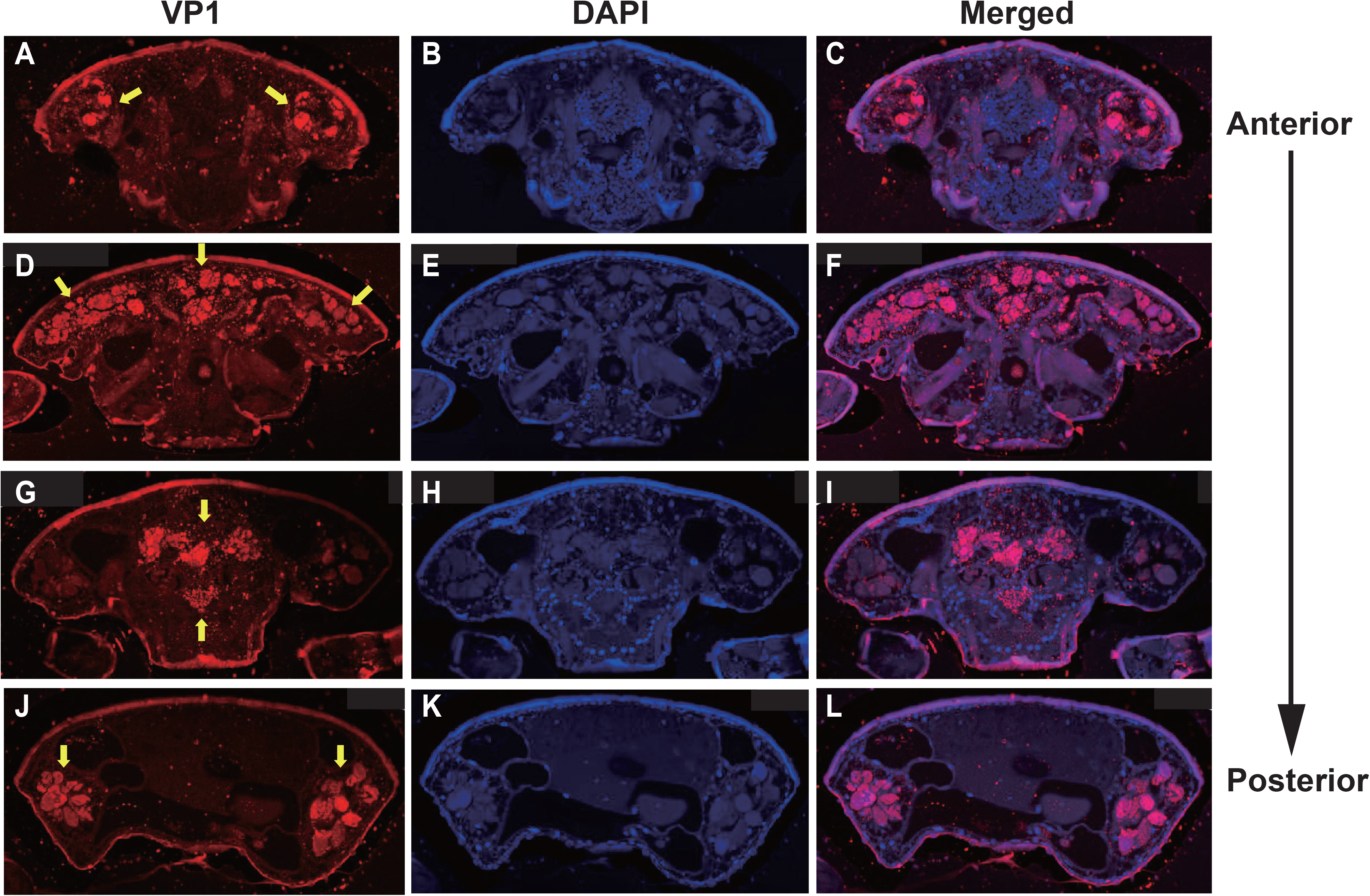
Localization of DWV in *T. mercedesae*. Transverse sections of *T. mercedesae* were immunostained by anti-VP1 antibody (VP1: A, D, G, and J) as well as DAPI (B, E, H, and K). The merged images (Merged: C, F, I, and L) are also shown. VP1-positive large dense spheres are indicated by yellow arrows. The dorsal side of mite is up and the anterior to posterior direction of mite body is also shown.

### *Hymenoptaecin* and *Defensin-1* mRNAs are induced in honey bee pupae by infestation of *T. mercedesae*

To understand the effects of *T. mercedesae* infestation on honey bees, we quantified mRNAs of two antimicrobial peptides (AMPs), *Hymenoptaecin* and *Defensin-1*, in the above pupae artificially infested with *T. mercedesae* and the control (uninfested) pupae. As shown in Fig. 4A and B, both *Hymenoptaecin* and *Defensin-1* mRNAs were increased by mite infestation. However, there was no significant correlation between the amounts of *Hymenoptaecin* and *Defensin-1* mRNAs expressed in individual mite infested pupae (*r* = −0.08, *P* < 0.78; Fig. 4C), suggesting that these AMPs were induced by different mechanisms during mite infestation. We then tested the correlation of DWV copy numbers and the amounts of either *Hymenoptaecin* or *Defensin-1* mRNA in both control and test pupae. The amount of *Defensin-1*, but not *Hymenoptaecin* (*r* = 0.141, *P* < 0.45) mRNA, was positively correlated with DWV copy number (Fig. 4D), suggesting that *Defensin-1* is induced by DWV infection and replication. Since *V. destructor* ingests the fat body cells of honey bees (Ramsey *et al.*, 2019), we first measured the relative amounts of honey bee *18S rRNA* in the individual mites to determine the degree of feeding of honey bee cells. A positive correlation was evident between the ingestion of honey bee cells by the mite and the amount of *Hymenoptaecin* mRNA, but there was no correlation for *Defensin-1* mRNA (*r* = 0.4648, *P* < 0.83; Fig. 4E).

**Figure 4.**
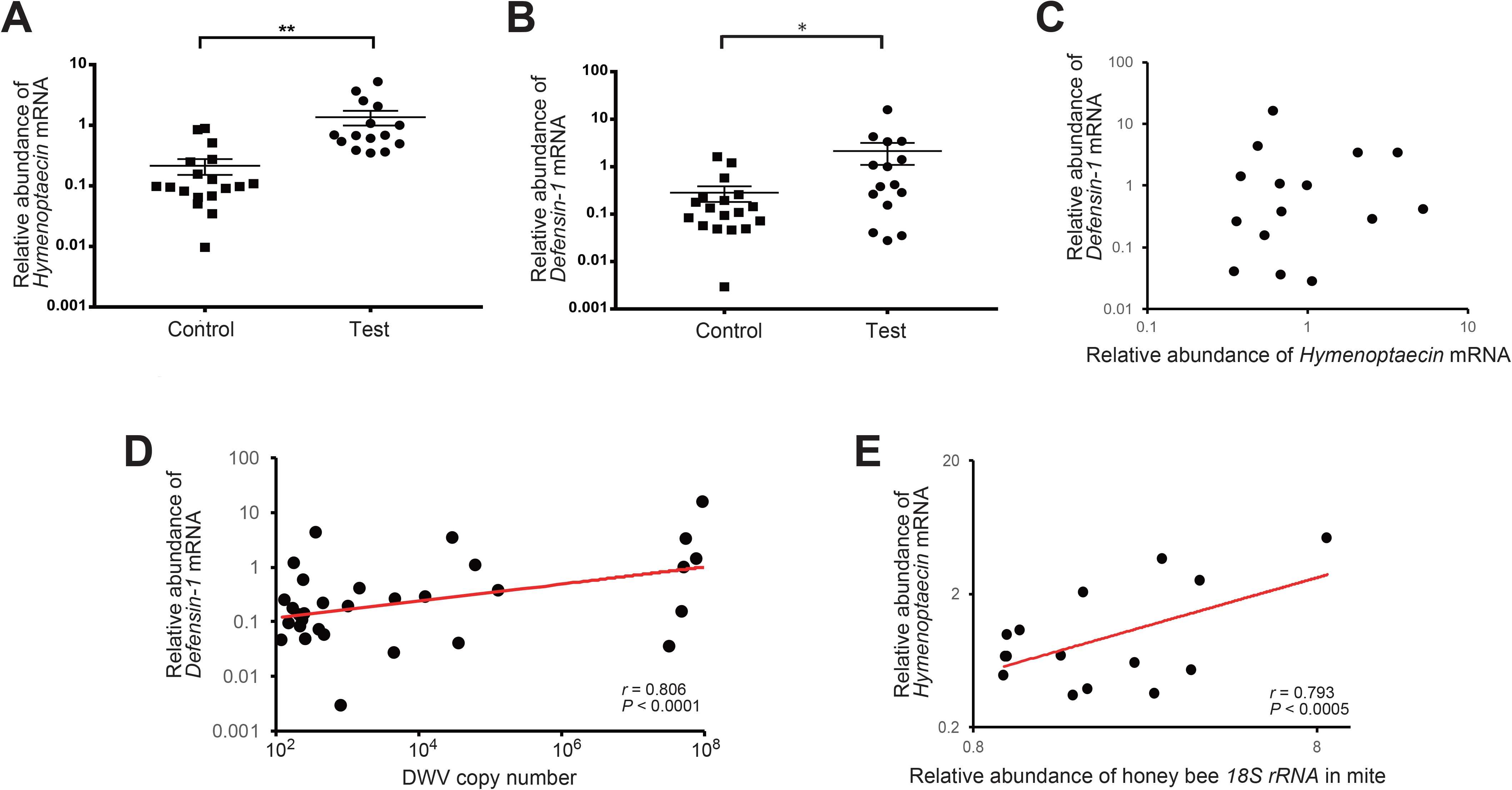
Increase of *Hymenoptaecin* and *Defensin-1* mRNAs by *T. mercedesae* infestation. Relative amounts of *Hymenoptaecin* (A) and *Defensin-1* (B) mRNAs in honey bee pupae artificially infested by *T. mercedesae* (Test, n =15) and the ones without mites (Control, n = 18). Mean values ± SEM (error bars) are shown. Brunner-Munzel test was used for the statistical analysis (*Hymenoptaecin*: **, *P* < 3.1E-08; *Defensin-1:* *, *P* < 0.042). (C) Relative amounts of *Hymenoptaecin* and *Defensin-1* mRNAs in individual honey bee pupae are plotted on the X- and Y-axis, respectively. (D) DWV copy number and relative amount of *Defensin-1* mRNA in individual honey bee pupae are plotted on the X- and Y-axis, respectively. The red line represents a fitted curve and the Pearson correlation value and *P*-value are also shown. (E) Relative amounts of *Hymenoptaecin* mRNA in individual honey bee pupae and honey bee *18S rRNA* in the infesting mites are plotted on the Y- and X-axis, respectively. The red line represents a fitted curve and the Pearson correlation value and *P*-value are also shown.

### Hymenoptaecin down-regulates *T. mercedesae Vg* gene

Since immune effector molecules, such as AMPs, are induced in honey bee pupae by *Tropilaelaps* mite infestation, the molecules could affect the physiology of the pupa, but also the mite, by feeding on the fat body and other tissues. *Vg* mRNA decreases with high load of DWV in *Tropilaelaps* mite by the transcriptome analysis (Supplementary Table 3). Five *Vg* related genes were identified in *T. mercedesae* genome (Accession numbers: OQR67440.1, OQR68606.1, OQR72029.1, OQR72561.1, and OQR79705.1). Based on the phylogenetic tree of *V. destructor* and *T. mercedesae* Vgs, OQR72561.1, OQR68606.1, and OQR79705.1 appeared to be *T. mercedesae* orthologs of *V. destructor* Vg-1, Vg-2, and XP_022657753.1, respectively. OQR72029.1 and OQR67440.1 were related to *V. destructor* Vg-1 and Vg-2, respectively (Supplementary Fig. 3). We tested the correlation between the amounts of the five *Vg* mRNAs in *T. mercedesae* and either *Hymenoptaecin* or *Defensin-1* mRNA in mite infested pupae. There was no correlation between *Defensin-1* mRNA and any of the five *Vg* mRNAs (Supplementary Table 4). However, a negative correlation was evident between *Hymenoptaecin* mRNA and *TmVg-1* or *TmVg-2*, but not between other *Vg* mRNAs (Fig. 5A, B, and Supplementary Table 4). These results suggest that expression of *T. mercedesae Vg* genes is downregulated by honey bee Hymenoptaecin. To test this hypothesis, we first established *Drosophila melanogastar* S2 cells expressing honey bee *Hymenoptaecin* (Supplementary Fig. 4), and then fed *T. mercedesae* with the extracts of S2 cells with or without Hymenoptaecin as previously described for *V. destructor* (Cabrera, Shirk and Teal, 2017). The amounts of the five *Vg* mRNAs were measured in individual mites and then compared. The experiments were repeated twice and the percentages of fed/survived mites were 35 (mean) ± 7.1 (SD) % for both treatments. Consistent with the above results, *TmVg-2* mRNA was significantly reduced in mites fed with the extract of S2 cells expressing *Hymenoptaecin* (Fig. 5C).

**Figure 5.**
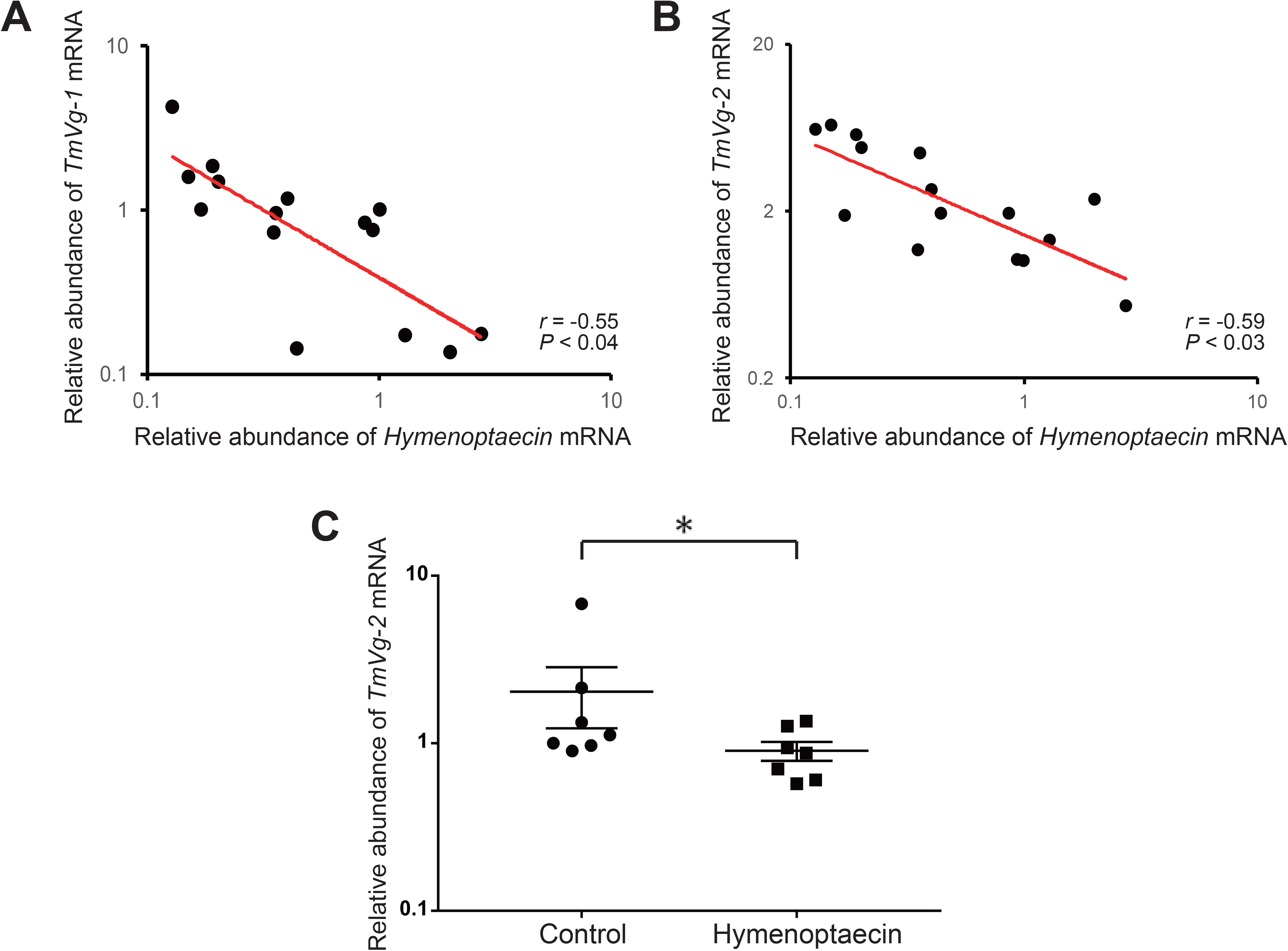
Hymenoptaecin down-regulates *T. mercedesae Vg* gene. Relative amounts of *Hymenoptaecin* mRNA in individual honey bee pupae and *TmVg-1* (A) and *TmVg-2* (B) in the infesting mites are plotted on the X- and X-axis, respectively. The red line represents a fitted curve and the Pearson correlation value and *P*-value are also shown. (C) Relative amount of *TmVg-2* mRNA in individual mites fed with the extracts of S2 cells expressing *Hymenoptaecin* or wild type S2 cells (Control). Mean values ± SEM (error bars) are shown. Brunner-Munzel test was used for the statistical analysis (*, *P* < 0.025).

## Discussion

### *T. mercedesae* functions as a vector for DWV without active replication inside mite

Our results show that the artificial infestation of honey bee pupae with *T. mercedesae* increases DWV copy number in pupae, but also transfers the variant present in the mite to the pupa. These results demonstrate that *T. mercedesae* functions as both mechanical and biological vectors by stimulating replication of the endogenous DWV in honey bee pupae and transmission of DWV. These properties are similar to those of *V. destructor* (Kuster, Boncristiani and Rueppell, 2014). Replication of the endogenous DWV can be induced by wounds caused by mite feeding. Consistent with previous reports for both *T. mercedesae* and *V. destructor* (Wu, Dong and Kadowaki, 2017; Posada-Florez *et al.*, 2019), the copy number of DWV could exceed 10^6^. However, we found that DWV had little effect on the transcriptome of *T. mercedesae.* Thus, DWV does not appear to significantly affect the fitness of mites. In fact, most DWV is present in the midgut lumen of mite, as previously reported with *V. destructor* (Zhang *et al.*, 2007; Santillan-Galicia *et al.*, 2008). Furthermore, RdRP was not detected in the mites with high DWV loads by western blotting. These results demonstrate that most of the DWV in the mite is derived from infested honey bee pupae and does not actively replicate inside the mite cells. Furthermore, the large difference in codon usage between DWV and *T. mercedesae* is unlikely to support the active translation of DWV proteins. Proteomic analysis of *T. mercedesae* and *V. destructor* failed to detect the non-structural proteins of honey bee viruses (Erban *et al.*, 2015; Dong *et al.*, 2017). Previous studies have suggested that the presence of a negative strand of DWV genome RNA in *V. destructor* is evidence of viral replication (Ongus *et al.*, 2004; Yue and Genersch, 2005).

However, this was recently questioned (Posada-Florez *et al.*, 2019). Therefore, our results do not support that the mites actively amplify the specific variant of DWV. The six pupa-mite pairs (Bee/Mite-3, 5, 10, 11, 13, and 15) contained four variants originally derived from the mites, indicating that they were specifically amplified in the honey bee over the endogenous variants. These results suggest that there is a mechanism to amplify the specific variant introduced by mite in honey bee. Since *T. mercedesae* is capable of transferring the associated variant to honey bee pupae, DWV must be present in the mite salivary gland. Although we did not detect DWV in the salivary gland by immunostaining of the sections, DWV has been previously detected in the salivary gland of *V. destructor* using mass spectrometry (Zhang and Han, 2019). The migration mechanism of DWV from the midgut to the salivary gland in the mites could be similar to that of mosquito-borne viruses, such as dengue virus and Zika virus in mosquitos (Cui *et al.*, 2019). It is also possible that selection of DWV variants may occur during migration from the midgut to the salivary gland. The mechanism of DWV transmission by honey bee mites as well as the virus/mite relationship could also help to understand the mechanisms of tick-borne viral diseases, for example, Tick-borne meningoencephalitis (Mansfield *et al.*, 2009), and mosquito-borne virus diseases.

### Downregulation of honey bee mite reproduction by host immune effector

Consistent with previous reports with *V. destructor* (Gregorc et al., 2012; (Kuster, Boncristiani and Rueppell, 2014), we found that *Defensin-1* and *Hymenoptaecin* mRNAs were induced by *Tropilaelapse* mite infestation, but by different mechanisms. Our results demonstrate that *Defensin-1* and *Hymenoptaecin* are induced by DWV replication and mite feeding, respectively. *Defensin-1* and *Hymenoptaecin* were reported to be under the control of the Toll and Imd pathways, respectively (Schlüns and Crozier, 2007; Lourenço *et al.*, 2018). These immune signaling pathways would be independently activated by the above events. We also demonstrated a negative correlation between *Hymenoptaecin* mRNA and *TmVg-1* or *TmVg-2* mRNA, and further demonstrated that Hymenoptaecin in S2 cell extract down-regulates expression of *TmVg-2* mRNA. Vg is a precursor for the major yolk protein vitellin, providing essential nutrients for the embryo (Tufail and Takeda, 2008). Thus, Vg and Vitellogenin receptor are essential for the reproduction of mites, such as *Panonychus citri* (Ali *et al.*, 2017). Accordingly, in *V. destructor*, high levels of *VdVg-1* and *VdVg-2* mRNAs are present at the oviposition stage (Cabrera Cordon *et al.*, 2013; McAfee *et al.*, 2017). A previous study also reported a negative association between the reproductive capability of *V. destructor* and honey bee immune gene mRNAs, such as *Relish*, *PGRP-S1*, and *Hymenoptaecin* (Kuster, Boncristiani and Rueppell, 2014). Hymenoptaecin is an AMP that was originally identified as a small positively-charged peptide capable of killing microorganisms by targeting the negatively charged membranes (Casteels *et al.*, 1993). However, the physiological roles of AMPs were not precisely characterized, and recent studies have revealed that they are also critical for tumor elimination, brain function, neurodegeneration, and aging (Hanson and Lemaitre, 2019). Since the Imd pathway is involved in processes that include viral defense, resistance to dessication, resistance to oxygen stress, and autophagy (Zhai, Huang and Yin, 2018), AMPs as well as the other downstream effectors could be involved in these processes. Thus, it is possible that Hymenoptaecin affects the mite tissue(s) to decrease the amount of *Vg* mRNA. Nevertheless, the mechanism of downregulation as well as the tissues synthesizing Vg remains to be elucidated. Hymenoptaecin can modulate specific cell signaling pathways in mite cells by indirect activation of receptors by displacing ligands, altering membrane microdomains, or directly acting as an alternate ligand. Given the present observation that Hymenoptaecin expressed in S2 cells failed to repress *TmVg-1* mRNA, it could be under the control of other downstream effectors of the Imd pathway. Hymenoptaecin induced by mite feeding could exert negative feedback on mite reproduction by repressing Vg synthesis. In fact, fecundity of both *T. mercedesae* and *V. destructor* is low (< 3 and < 7, respectively) and significant fractions of non-reproductive mites have also been described (Nazzi and Le Conte, 2016; Chantawannakul *et al.*, 2018). This equilibrium between the host (honey bee) and parasite (mite) could be established by the interaction between Hymenoptaecin and mite Vg. Our study should help further explorations of the novel physiological roles of AMPs in honey bees and other insects.

## Author contribution

TK conceived and designed research strategy. YW, QL, and BW performed research. YW, MK, and TK analyzed data. YW, MK, and TK drafted and edited the paper.

## Acknowledgements

We thank Gongjie Wu, Nihao Sun, and Meng Yuan for their contribution to conduct the experiments. We also thank Dr. Ferdinand Kappes for his help to prepare the mite sections.

## Materials and Methods

### Artificial infestation of honey bee pupae with mites

*A. mellifera* colonies were obtained from local beekeepers and maintained at Xi’an Jiaotong-Liverpool University. Honey bee pupae (n = 33) with white eyes were sampled from the mite-free colony by opening the capped brood cells. Adult female mites (n = 15) were collected from another colony heavily infested with *T. mercedesae* as above. A single pupa and mite were put inside a gelatin capsule. As the control, the remaining pupae (n = 18) were individually incubated without the mite. The capsules were inserted to a tube rack vertically positioned in an incubator at 33 °C with 70 % relative humidity for a week (Egekwu *et al.*, 2018).

### Isolation of total RNA and RT-PCR

Head was first dissected from each pupa and total RNA was extracted from the individual pupal heads and mites using TRI Reagent® (Sigma-Aldrich) according to the manufacturer’s instruction. Glycogen (1 μg) was added to facilitate isopropanol precipitation of the mite RNA sample. Reverse transcription (RT) reaction was carried out using 1 μL of total RNA, random primer (TOYOBO), ReverTra Ace (TOYOBO), and RNase inhibitor (Beyotime). RNase H (Beyotime) was then added to digest RNA in RNA/cDNA heteroduplex after cDNA synthesis. DWV in the honey bee and mite samples was detected by RT-PCR using DWV #1 primers (Supplementary Table 5) and the cycling condition of 2 min at 94 ° followed by 32 cycles of 10 sec at 98 °, 20 sec at 55 °, and 30 sec at 68 °. The PCR products were analysed by 2 % agarose gel. *A. mellifera* and *T. mercedesae EF-1α* mRNAs (Supplementary Table 5) were used as the positive controls to verify successful RT.

### Sequencing of RT-PCR products

PCR products obtained by above RT-PCR were purified by QIAquick PCR Purification Kit (QIAGEN) according to the manufacturer’s instruction, and then directly sequenced by Sanger method using the PCR primers.

### Analysis of DWV, honey bee *AMP*, and mite *Vg* mRNAs by qRT-PCR

DWV copy number was determined by qRT-PCR using a Hieff™® qRT-PCR SYBR Green Master Mix (Low Rox Plus, Yesen) and DWV #2 primers (Supplementary Table 5). To prepare a standard curve for DWV, PCR product obtained by above primers was purified and the copy number was determined by a formula below.

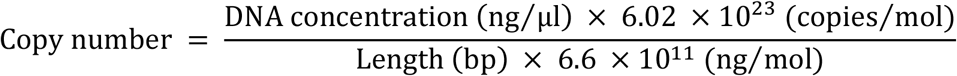

6.6 × 10^11^ng/mol is the average molecular mass of one base pair and 6.02 × 10^23^ copies/mol is Avogadro’s number. We conducted qPCR using 10^1^-10^9^ copy number of the PCR product and then plotted the Ct values against the log values of copy numbers. DWV copy number in the sample was determined using the standard curve. The amount of cDNA added to each qPCR reaction was normalized using either *A. mellifera* or *T. mercedesae 18S rRNA* as the endogenous reference (Supplementary Table 5).

The relative amounts of *Hymenoptaecin*, *Defensin-1*, and *T. mercedesae Vg* mRNAs in the samples were measured by the ΔΔCt method. The primers for *Hymenoptaecin*, *Defensin-1*, and five *T. mercedesae Vgs* are listed in Supplementary Table 5. *A. mellifera* or *T. mercedesae 18S rRNA* was used as the endogenous reference.

### Effects of introducing wound on DWV copy numbers in honey bee pupae

The mite-free honey bee pupae with pink eyes were sampled from *T. mercedesae-*infested colony. A small physical wound was introduced to the thorax (n = 6) with a sterilized microliter syringe (GAOGE) and control pupae (n = 7) were untreated. All pupae were then individually put in a gelatin capsule and incubated for 37 hours and DWV copy numbers in the individual pupal heads were measured as above.

### Raising antibodies against VP1 and RdRP of DWV

The P-domain of VP1 (amino acid 748-901 of DWV polyprotein) was PCR amplified using 5’-Nde I-P-domain and 3’-Xho I-P-domain primers (Supplementary Table 5). The PCR amplicon was digested with Nde I (NEB) and Xho I (NEB). The part of RdRP (amino acid 2563-2797 of DWV polyprotein) was also PCR amplified using 5’-Kpn I-RdRP and 3’-Hind III-RdRP primers (Supplementary Table 5). The PCR product was digested with Kpn I (NEB) and Hind III (NEB). The restriction enzyme digested DNA fragments were purified and then cloned to the corresponding sites in pCold-I vector (TAKARA) followed by transformation to *E. coli* BL21.

The transformed *E. coli* was grown in LB medium containing ampillicin (0.1 mg/ml) at 37 °C until the A_600_ reached to approximate 0.5, and then the culture was cooled down and added with isopropyl-thio-galactoside at the final concentration of 0.5 mM to induce the protein expression at 15 °C overnight. *E. coli* was suspended with 100 mL of lysis buffer (50 mM NaH_2_PO_4_, 300 mM NaCl, 1 mM DTT, and protease inhibitors at pH 8.0). The cell suspension was then sonicated for 45 min (30 sec sonication with 3 min interval) at amplitude 100 on ice using Q700 sonicator (Qsonica). After centrifugation, the supernatant was collected and incubated with His-tag Protein Purification Resin (Beyotime) at 4°C for 2 hours. The resin was then washed five times with 10 mL of washing buffer (50 mM NaH_2_PO_4_, 300 mM NaCl, and 2 mM imidazole, pH 8.0), and then the bound protein was eluted six times with 1 mL then twice with 5 mL of elution buffer (50 mM NaH_2_PO_4_, 500 mM NaCl, and 250 mM imidazole, pH 8.0). The purified proteins were dialyzed against PBS at 4°C overnight, and then concentrated using Vivaspin^®^ 6 polyethersulfone 10 kDa (Sartorius). Concentrations of the purified proteins were measured using BCA protein assay kit (Beyotime) according to the manufacturer’s instruction. Purification of the RdRP peptide was carried out using above buffers containing 0.1 % sarcosyl to increase the protein solubility. The purified proteins were delivered to a company (GeneScript-Nanjing) to obtain the affinity purified rabbit polyclonal antibodies.

### SDS-PAGE and western blot

The proteins were suspended with the sample buffer (2 % SDS, 10 % glycerol, 10 % β-mercaptoethanol, 0.25 % bromophenol blue, and 50 mM Tris-HCl, pH 6.8), and then heated at 99 °C for 5 min followed by applying to 12 % SDS-PAGE gel. After electrophoresis, the gel was treated with the staining buffer (0.25 % Coomassie brilliant blue G-250, 40 % methanol, and 10 % acetic acid), and then washed with the destaining buffer (40 % methanol and 10 % acetic acid). The bands were visualized and analyzed using ChemiDoc^™^ MP imaging system and Image Lab^™^ touch software (BIO-RAD).

Honey bee pupal heads and *Tropilaelaps* mites were individually homogenized with 300 and 50 μL of sample buffer, respectively. *Drosophila* S2 cells in 12-well plate were lysed with 200 μL of sample buffer. All samples were heated as above and centrifuged, and then the supernatants were applied to 10 (for honey bee and mite lysates) or 15 % (for S2 cell lysates) SDS-PAGE gel. After electrophoresis, proteins in the gel were transferred to a pure nitrocellulose blotting membrane (Pall^®^ Life Sciences). The membrane with proteins was first blocked with PBST (0.1 % Tween-20/PBS) containing 5 % BSA at room temperature for 1 hour, and then incubated with 1000-fold diluted primary antibody (anti-VP1 antibody, anti-RdRP antibody, or anti-His tag antibody) in above buffer at 4 °C overnight. The membrane was washed with PBST three times for 5 min each, and then incubated with 10,000-fold diluted IRDye® 680RD donkey anti-rabbit IgG (H+L) (LI-COR Biosciences) in PBST containing 5 % skim milk at room temperature for 1.5 hours. The membrane was washed as above, and then visualized using Odyssey Imaging System (LI-COR Biosciences).

### Analysis of mite transcriptomes by RNA-seq

Individual *Tropilaelaps* mites were tested for DWV by RT-PCR as above and separated to two groups each for high (High_A and High_B) and low (Low_A and Low_B) DWV levels. Each group was made of total RNAs prepared from ten mites and sequenced at BGI (Shenzhen, China) using Illumina HiSeq 4000 platform. After sequencing, the raw data were filtered to remove the adaptor sequences, contamination, and low-quality reads by BGI. The Quality control (QC) was further analyzed using FastQC. All RNA-seq data are available in SRA database with the accession #: SUB6997575.

The reference genome and annotated genes of *T. mercedesae* were first acquired from NCBI (https://www.ncbi.nlm.nih.gov/genome/53919?genome_assembly_id=313451), and then used for building the index by Hisat2–build indexer (Kim, Langmead and Salzberg, 2015). The generated index files were used to align the clean reads of four RNA-seq samples to the reference genome. Subsequently, SAM file outputs from the previous step were sorted using SAMtools (Li *et al.*, 2009). HTSeq-count (Anders, Pyl and Huber, 2015) was further applied to obtain the raw read counts for downstream analysis of identifying the differentially expressed genes (DEGs) between the mites with low and high DWV loads in *R* (V3.4.3) based Bioconductor edgeR package (V3.20.9) (Robinson, McCarthy and Smyth, 2010). The DEGs were cut-off by a False Discovery Rate (FDR) at 0.05

### Immunostaining of the mite thin sections

*Tropilaelaps* mites collected from the hive were washed with bleach and PBS, and then fixed with 4 % paraformaldehyde/PBS at 4°C for 24 hours with gentle shaking. The fixed samples were stored in methanol at −20 °C and then converted into absolute n-butanol followed by embedding in Technovit 8100 (Kulzer, Wehrheim, Germany). On the rotation microtome (Leica RM 2245), 8 μm thick serial sections were prepared with glass knives and placed into water drops on silanized slides. The sections were dried for 30 minutes at 50 °C. The above thin sections were washed five times with PBS, and then treated with 2 mg/mL pepsin in 0.9 % NaCl, pH 2.0 at 37 °C for 10 min. They were then washed three times with PBS, three times with PT (PBS containing 0.1 % Triton-X 100), and once with PBS. The sections were blocked with PBS containing 3 % BSA and 1 % normal goat serum at 4 °C overnight and then they were alternatively incubated with 1,000-fold diluted anti-VP1 antibody and the pre-immune serum in above buffer at 4 °C overnight. After washing eight times with PBS, the sections were incubated with 1,000-fold diluted Goat anti-rabbit IgG (H+L) Superclonal™ secondary antibody, Alexa Fluor 555 (Thermo Scientific) at room temperature for 1.5 hours. After washing seven times with PBS, the sections were incubated with 0.5 μg/mL DAPI (Beyotime) at room temperature for 15 min. Following the final wash with PBS, the sections were mounted with Antifade mounting medium (Beyotime). Each wash was conducted for 10 min. The immunostained sections above were observed using a confocal microscope, LSM 880 (Zeiss) with the TileScan method. ImageJ was used for the image analysis.

### Phylogenetic trees of the DWV isolates and mite Vgs

The representative RNA sequences of DWV isolates from the honey bee pupae and mites were aligned using MUSCLE (Edgar, 2004), and then Tamura 3-parameter+G was selected as the best-fit substitution model for constructing the phylogeny. The condensed phylogenetic tree was constructed using the maximum likelihood method and a bootstrap value of 1,000 replicates with MEGA7 (Kumar, Stecher and Tamura, 2016). Amino acid sequences of all *T. mercedesae* and *V. destructor* Vgs were retrieved from NCBI and the phylogenetic tree was constructed as above except Jones-Taylor-Thornton+G+F was used as the best model.

### Codon usage analysis

Codon usage tables to show the codon frequencies per 1000 codons for DWV, *A. mellifera*, and *T. mercedesae* were obtained using HIVE-CUTs (Athey *et al.*, 2017). To compare the codon frequencies between DWV and either *A. mellifera* or *T. mercedesae*, three stop codons, and codons for methionine and tryptophan were omitted from the analysis.

### Establishing *Drosophila* S2 cells to stably express Hymenoptaecin

*Hymenoptaecin* cDNA was obtained by RT-PCR using honey bee RT and Hymenoptaecin #1 primers (Supplementary Table 5). The PCR amplicon was digested with EcoR I and Age I and cloned to the same sites in pAc5.1/V5-His B (Thermo Scientific) to express the His-tagged protein. S2 cells in 12-well plate were transfected with either 2 μg of Hymenoptaecin expression vector or empty vector and 10 μL of HilyMax (DOJINDO) for 24 hours. After replacing the medium, the cells were cultured for 24 hours, and then analyzed by western blot using rabbit polyclonal anti-His tag antibody (BBI). After confirming the protein expression, the untagged version of *Hymenoptaecin* cDNA was PCR amplified using Hymenoptaecin #2 primers (Supplementary Table 5) and digested with Xho I and EcoR V followed by inserting to the same sites in pMK33/pMtHy (Koelle *et al.*, 1991). S2 cells were transfected with this expression vector or empty pMK33/pMtHy as above and the stable transfectants were first selected by 0.1 mg/mL hygromycin for 10 days, and then the concentration was increased to 0.3 mg/mL. The hygromycin-resistant S2 cells (2 × 10^7^) were harvested during the logarithmic growth phase and suspended with 0.3 mL of Grace medium followed by sonication and store at −20 °C.

### Feeding mites with S2 cell extracts

Feeding *Tropilaelaps* mites was carried out in an inverted flat-cap 2 mL microcentrifuge tube. The bottom of the tube was cut and plugged with cotton to provide ventilation and the cap contained a small piece of a sterile cotton ball. S2 cell extracts prepared as above together with 2 % Royal blue (50 μL) were dispensed onto the cotton and then 10 starved mites were added prior to closing the tube. The assay tubes were placed in an incubator at 33 °C with 90 % relative humidity for 24 hours under dark. The fed and survived mites were collected and then five *Vg* mRNAs were analysed by qRT-PCR.

---

**Supplementary Figure 1.**
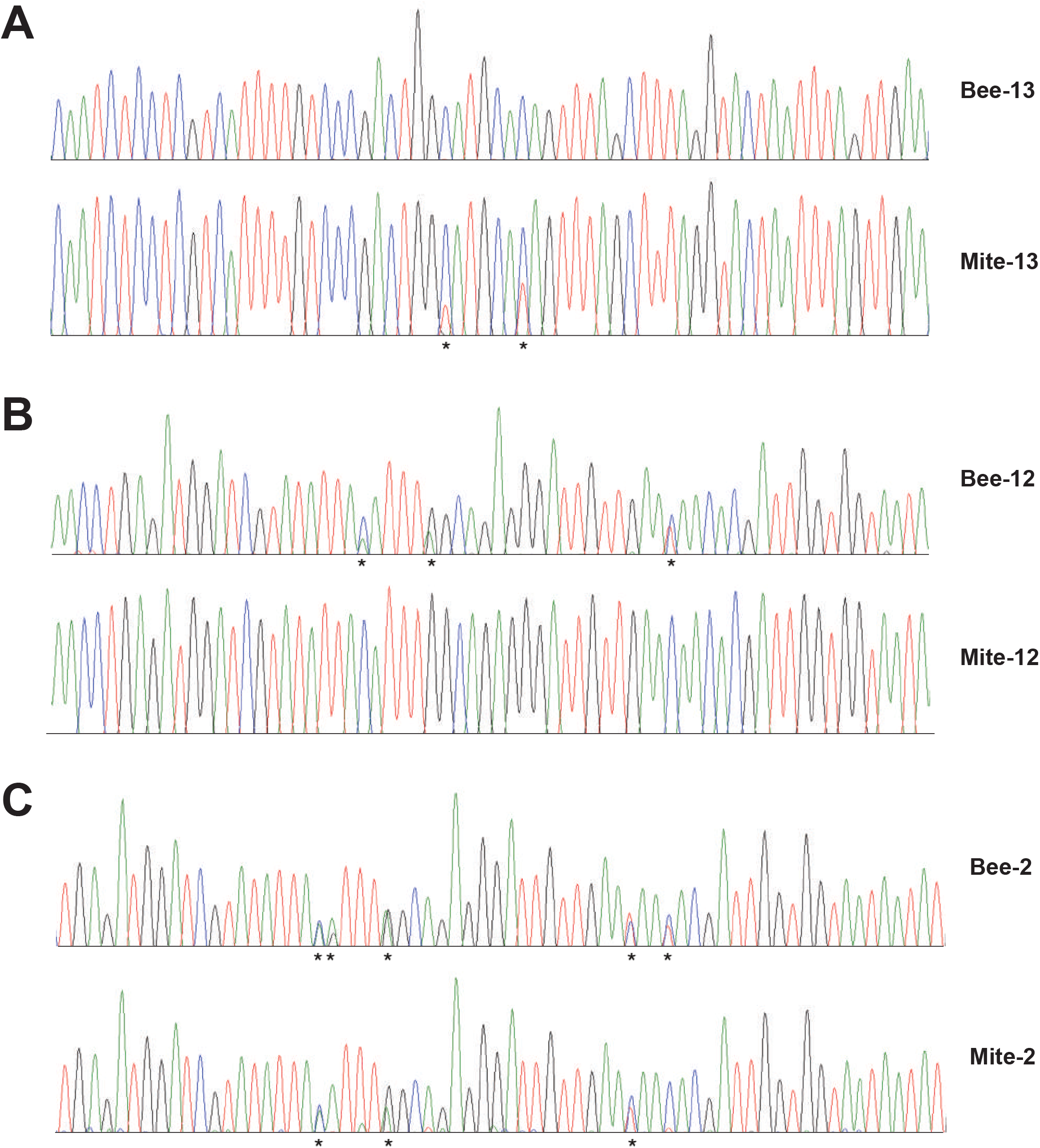
Presence of single and multiple DWV variants in honey bee pupae and the infesting *T. mercedesae*. Representative Sanger sequencing electropherograms of the RT-PCR products show (A) the presence of single variant in pupa and multiple variants in the infesting mite (Bee/Mite-13), (B) the presence of multiple variants in pupa and single variant in the infesting mite (Bee/Mite-12), and (C) the presence of multiple variants in both pupa and the infesting mite (Bee/Mite-2). Only single peaks are present at all positions for the single variant; however, two peaks (show by asterisks) are present at several positions for the multiple variant.

**Supplementary Figure 2.**
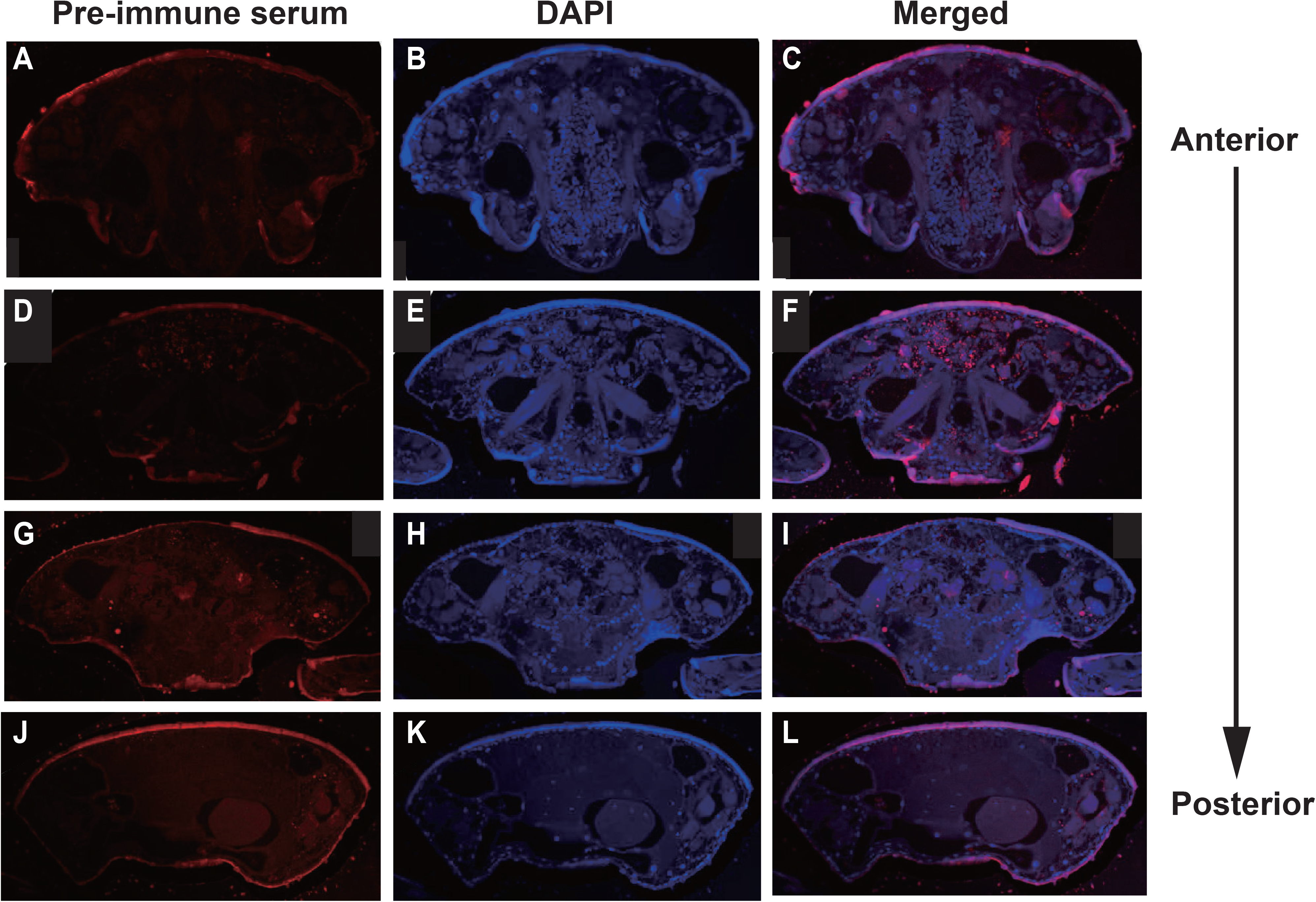
Immunostaining of *T. mercedesae* sections by the pre-immune serum. Transverse sections of *T. mercedesae* were immunostained by the pre-immune serum (A, D, G, and J) as well as DAPI (B, E, H, and K). The merged images (Merged: C, F, I, and L) are also shown. There is no specific staining in all sections. The anterior to posterior direction of mite body is also shown.

**Supplementary Figure 3.**
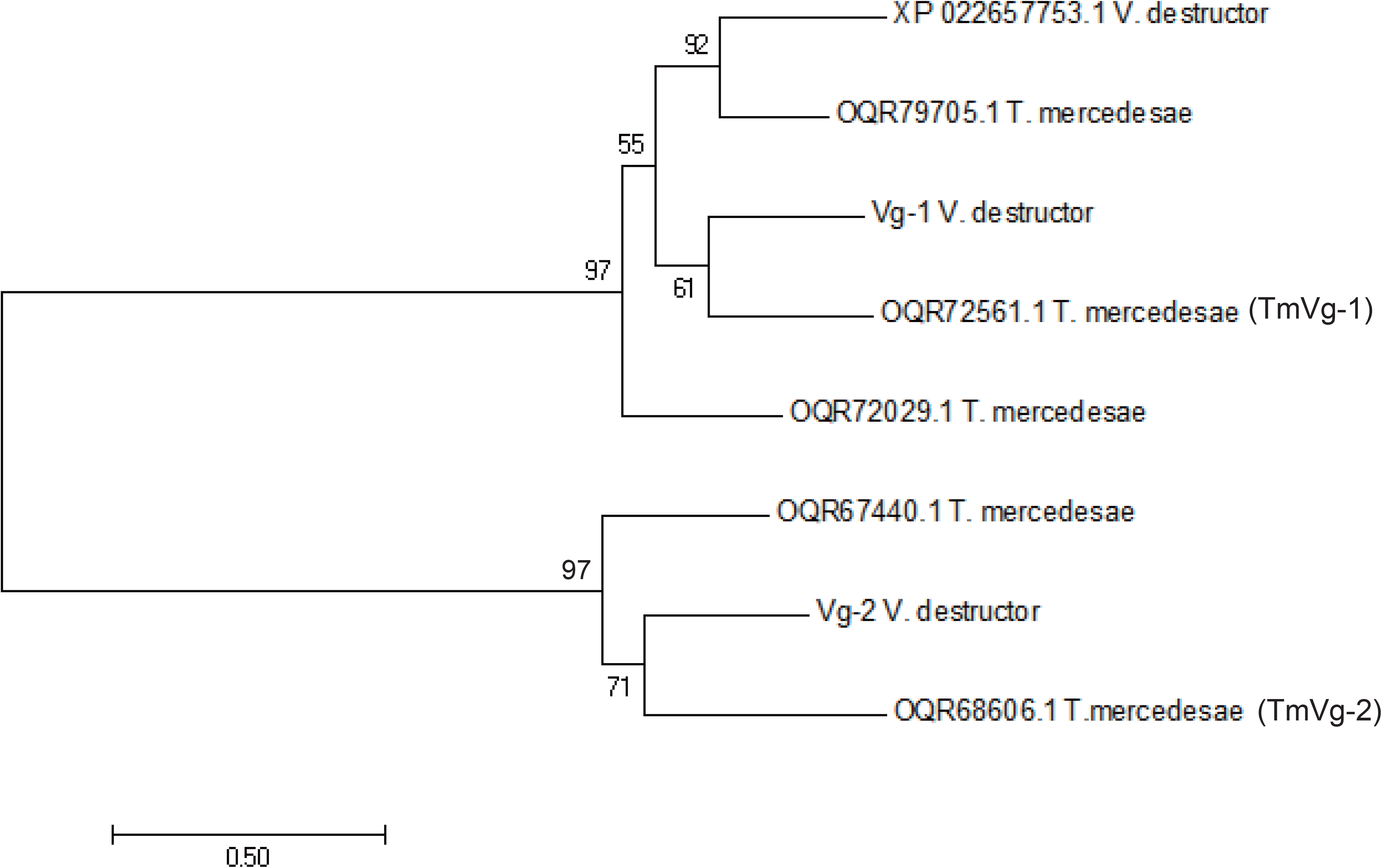
Phylogeny of *T. mercedesae* and *V. destructor* vitellogenins (Vgs) Phylogenetic tree of five and three Vgs of *T. mercedesae* and *V. destructor* was constructed by maximum-likelihood method. Bootstrap values are shown at the corresponding node of each branch.

**Supplementary Figure 4.**
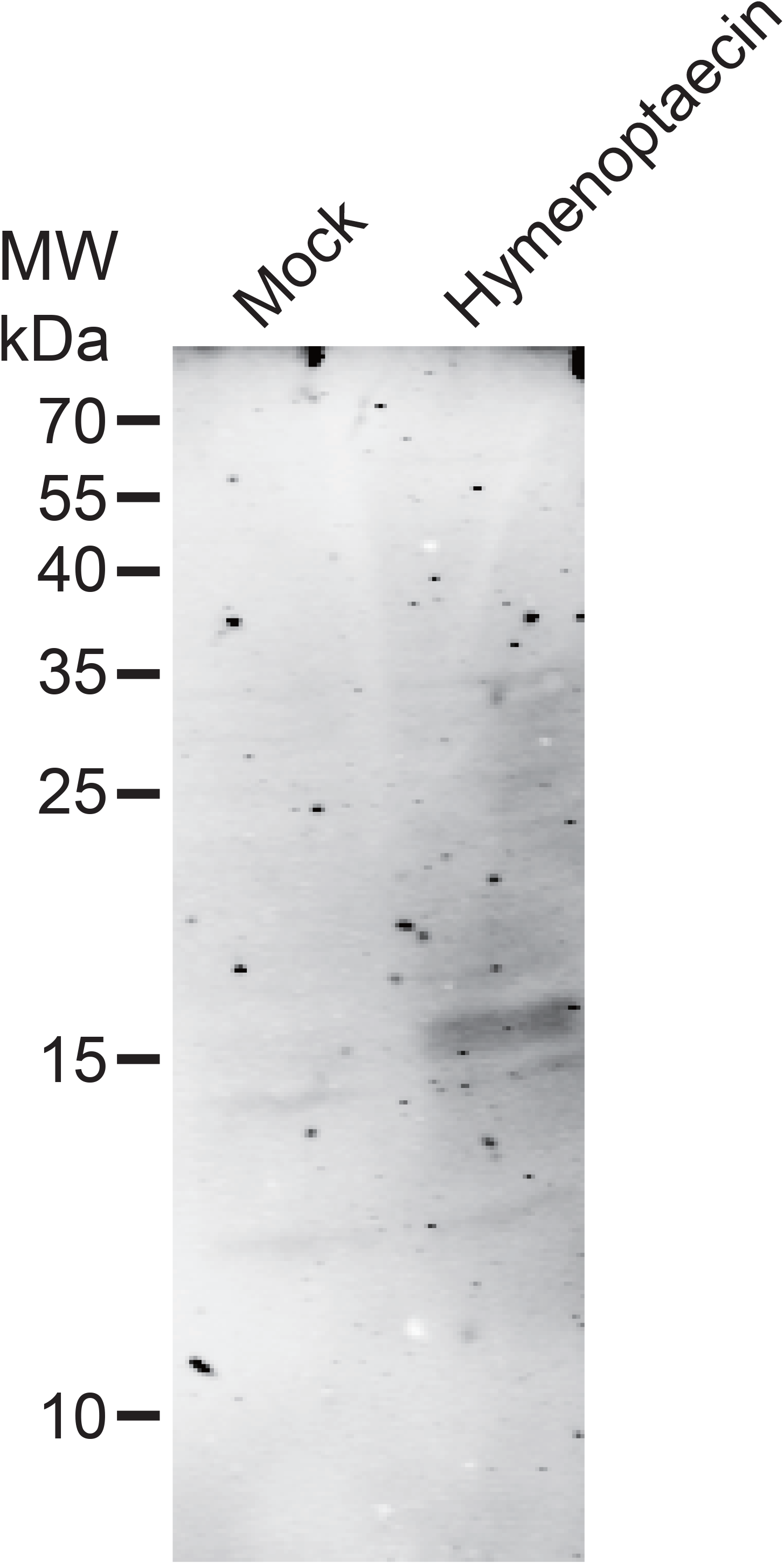
Expression of Hymenoptaecin in S2 cells. Lysates of S2 cells transfected with either empty vector (Mock) or *Hymenoptaecin*-expressing construct were analyzed by western blot using anti-His tag antibody. The size of protein molecular weight marker (MW) is at the left. A major 15kDa protein is present in *Hymenoptaecin*-expressing S2 cells.

**Supplementary Table 1.** Profile of DWV infection in honey bee pupae and the infesting *Tropilaelaps* mites.

|  | Single variant in pupa | Multiple variants in pupa |
| --- | --- | --- |
| Single variant in mite | 1, 3, 4, 7, 8, 9, 10, 11, 14, 15 | 12 |
| Multiple variants in mite | 6, 13 | 2, 5 |

The pupa/mite pairs in the clusters with high and low copy numbers of DWV in Figure 1 are indicated by red and black letters, respectively.

**Supplementary Table 2.** Alignment rates of RNA-seq reads to *T. mercedesae* and DWV genomes

|  | Alignment rate to *T. mercedesae* genome (%) | Alignment rate to DWV (%) |
| --- | --- | --- |
| High-A | 37.73 | 22.45 |
| High-B | 39.64 | 23.12 |
| Low-A | 80.44 | 0.08 |
| Low-B | 85.53 | 0.01 |

**Supplementary Table 3.** *T. mercedesae* genes up-regulated by DWV

| Gene | Annotation | P-value | FDR |
| --- | --- | --- | --- |
| OQR67497.1 | lipase member H-A-like<br>[*Tropilaelaps mercedesae*] | 1.49759E-06 | 0.00286 |
| OQR72005.1 | chymotrypsin elastase family member<br>3B-like [*Tropilaelaps mercedesae*] | 8.45038E-06 | 0.01005 |
| OQR70413.1 | hypothetical protein BIW11_04159<br>[*Tropilaelaps mercedesae*] | 3.43603E-05 | 0.03143 |
| OQR68960.1 | hypothetical protein BIW11_12564<br>[*Tropilaelaps mercedesae*] | 4.31267E-05 | 0.03663 |
| OQR77579.1 | hypothetical protein BIW11_06990,<br>partial [*Tropilaelaps mercedesae*] | 5.29712E-05 | 0.042 |

*T. mercedesae* genes down-regulated by DWV
| Gene | Annotation | P-value | FDR |
| --- | --- | --- | --- |
| OQR70206.1 | zinc finger protein-like<br>[Tropilaelaps mercedesae] | 1.7968E-14 | 2.1E-10 |
| OQR70191.1 | hypothetical protein BIW11_11788<br>[Tropilaelaps mercedesae] | 1.85428E-08 | 0.00011 |
| OQR74958.1 | hypothetical protein BIW11_00866<br>[Tropilaelaps mercedesae] | 1.68204E-06 | 0.00286 |
| OQR67746.1 | nose resistant to fluoxetine protein<br>6-like [Tropilaelaps mercedesae] | 7.87212E-07 | 0.00286 |
| OQR71056.1 | hypothetical protein BIW11_11234<br>[Tropilaelaps mercedesae] | 1.02906E-06 | 0.00286 |
| OQR68367.1 | hypothetical protein BIW11_12957<br>[Tropilaelaps mercedesae] | 1.22238E-06 | 0.00286 |
| OQR76942.1 | hypothetical protein BIW11_07447<br>[Tropilaelaps mercedesae] | 2.57456E-06 | 0.00383 |
| OQR75105.1 | cement protein RIM36-like<br>[Tropilaelaps mercedesae] | 6.00625E-06 | 0.00794 |
| OQR79705.1 | vitellogenin 1-like [Tropilaelaps<br>mercedesae] | 1.70535E-05 | 0.0169 |
| OQR67513.1 | Larval cuticle protein A3A-like<br>[Tropilaelaps mercedesae] | 1.69746E-05 | 0.0169 |

**Supplementary Table 4.** Pearson correlation analysis between either *Defensin-1* or *Hymenoptaecin* mRNA and five *Vitellogenin*-like mRNAs of *T. mercedesae*

|  | *TmVg-1* | *TmVg-2* | *OQR79705.1* | *OQR72029.1* | *OQR67440.1* |
| --- | --- | --- | --- | --- | --- |
| *Defensin-1* | $r = -0.12$,<br>$P < 0.67$ | $r = 0.08$,<br>$P < 0.77$ | $r = 0.15$,<br>$P < 0.59$ | $r = -0.1$,<br>$P < 0.74$ | $r = -0.14$,<br>$P < 0.61$ |
| *Hymenoptaecin* | $r = -0.55$,<br>$P < 0.04$ | $r = -0.59$,<br>$P < 0.03$ | $r = -0.44$,<br>$P < 0.1$ | $r = -0.51$,<br>$P < 0.06$ | $r = -0.41$,<br>$P < 0.14$ |

The correlation values and *P*-values are shown.

**Supplementary Table 5.**
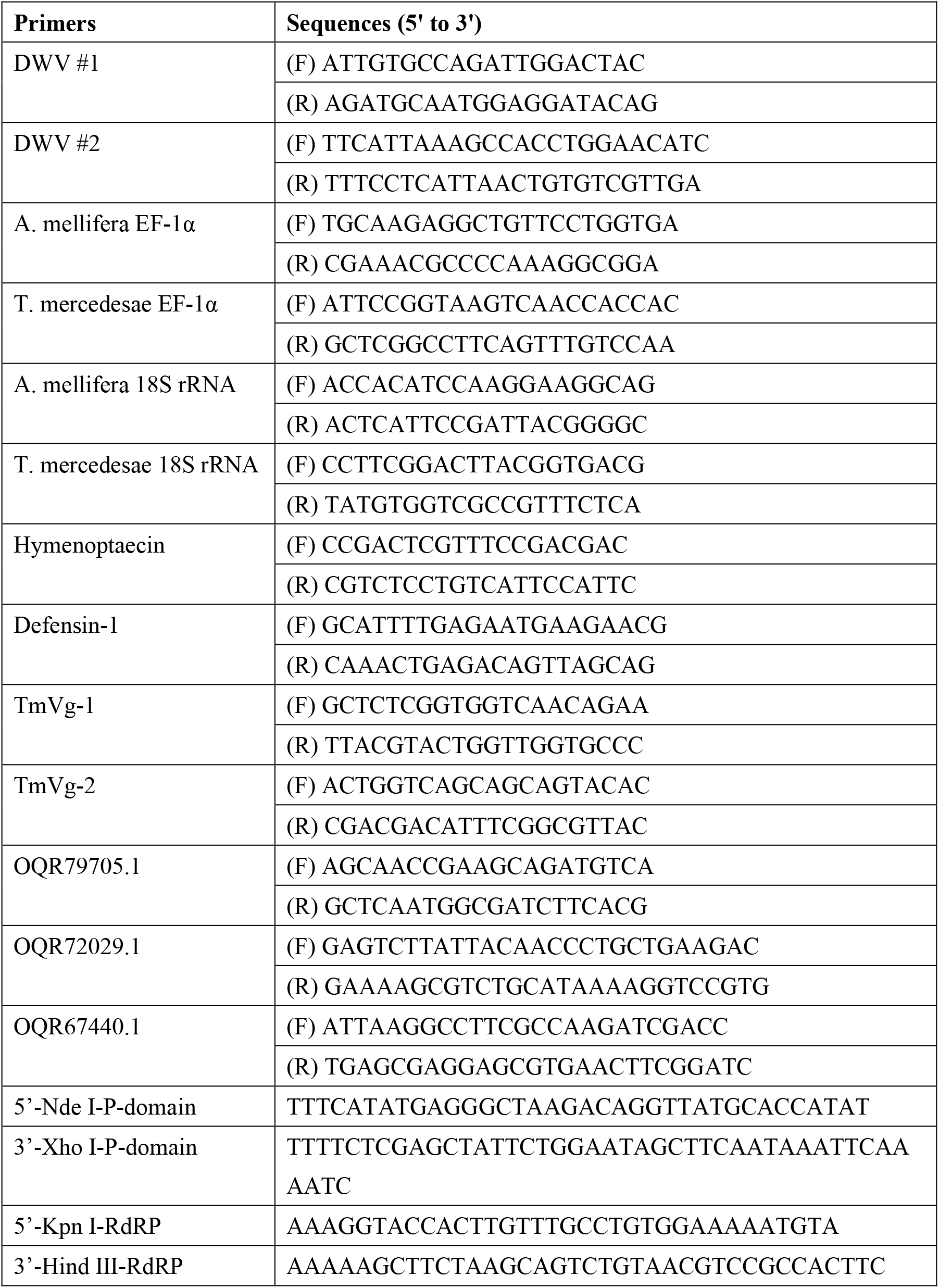
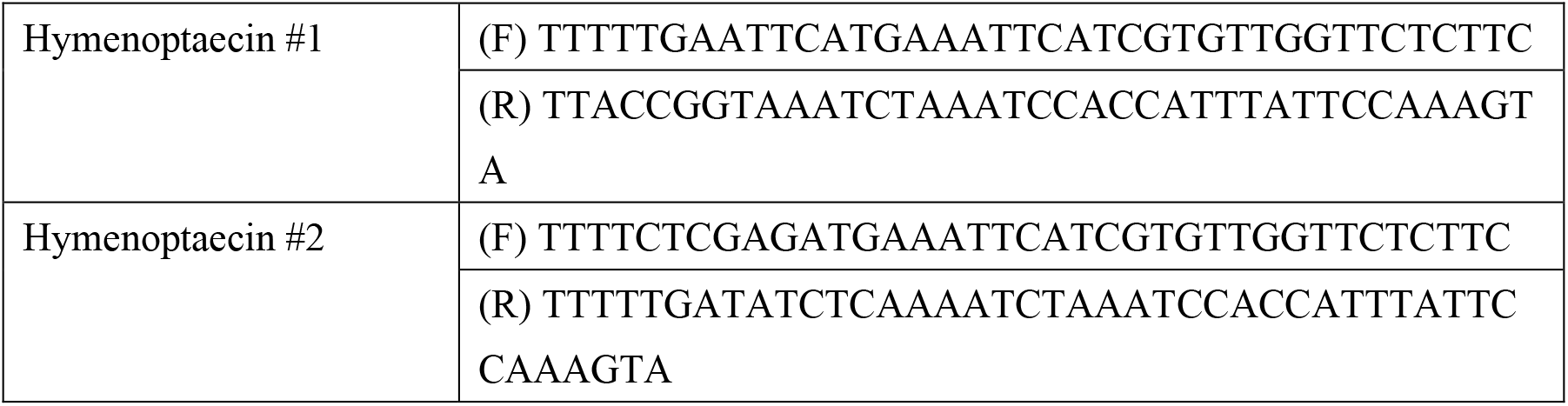
List of primers used in this study

